# A Synthetic Biology Approach to Sequential Stripe Patterning and Somitogenesis

**DOI:** 10.1101/825406

**Authors:** Fuqing Wu, Changhan He, Xin Fang, Javier Baez, Thai Ohnmacht, Qi Zhang, Xingwen Chen, Kyle R. Allison, Yang Kuang, Xiao Wang

## Abstract

Reaction-diffusion (RD) based clock and wavefront model has long been proposed as the mechanism underlying biological pattern formation of repeated and segmented structures including somitogenesis. However, systematic molecular level understanding of the mechanism remains elusive, largely due to the lack of suitable experimental systems to probe RD quantitatively *in vivo*. Here we design a synthetic gene circuit that couples gene expression regulation (reaction) with quorum sensing (diffusion) to guide bacterial cells self-organizing into stripe patterns at both microscopic and colony scales. An experimentally verified mathematical model confirms that these periodic spatial structures are emerged from the integration of oscillatory gene expression as the molecular clock and the outward expanding diffusions as the propagating wavefront. Furthermore, our paired model-experiment data illustrate that the RD-based patterning is sensitive to initial conditions and can be modulated by external inducers to generate diverse patterns, including multiple-stripe pattern, target-like pattern and ring patterns with reversed fluorescence. Powered by our synthetic biology setup, we also test different topologies of gene networks and show that network motifs enabling robust oscillations are foundations of sequential stripe pattern formation. These results verified close connections between gene network topology and resulting RD driven pattern formation, offering an engineering approach to help understand biological development.

## Main text

Turing’s seminal work first proposed reaction-diffusion (RD) as the "chemical basis of morphogenesis" over six decades ago ^1^. It provides a general theoretical foundation of pattern formation via RD mechanisms. Two decades later, RD driven clock and wavefront (CW) mechanism was hypothesized as the mechanism underlying formation of repeated and segmented structures such as somites in development ^2^. Since then, although RD driven pattern formation has been demonstrated or identified in chemical, physical, and ecological systems ^3–10^, its much-hypothesized role in multicellular pattern formation hasn’t been fully studied biologically. This is largely due to the lack of suitable model systems to test such hypotheses. For example, somite development requires precise temporal and spatial coordination between a heterogeneous web of intracellular responses and intercellular communications, both under control of complex gene regulation networks and influences of universal gene expression stochasticity. Such complexity poses a great challenge to fully understand mechanistic basis of somite formation *in vivo*. Engineered microbes carrying rationally designed gene circuits provide an effective venue to study this problem from bottom up. Previous studies using synthetic circuits have demonstrated formation of predefined patterns, cell motility based stripe formation, and scale invariant ring pattern formation ^11–15^. However, gene network directed RD based clock and wavefront pattern formation, despite its importance in developmental biology and extensive theoretical studies ^16–22^, has not been experimentally realized.

Past studies have suggested that nonlinear multistable systems could also direct spatiotemporal pattern formation when coupled with external diffusion process ^23–25^. Following this strategy to achieve a multicellular pattern formation, we designed and constructed a mutually inhibitory network with positive autoregulation and communications (MINPAC) by expanding our previously demonstrated quadrastable gene circuit ^26^ with added quorum-sensing modules to enable intercellular communications (Fig. 1A and 1B).

**Fig. 1.**
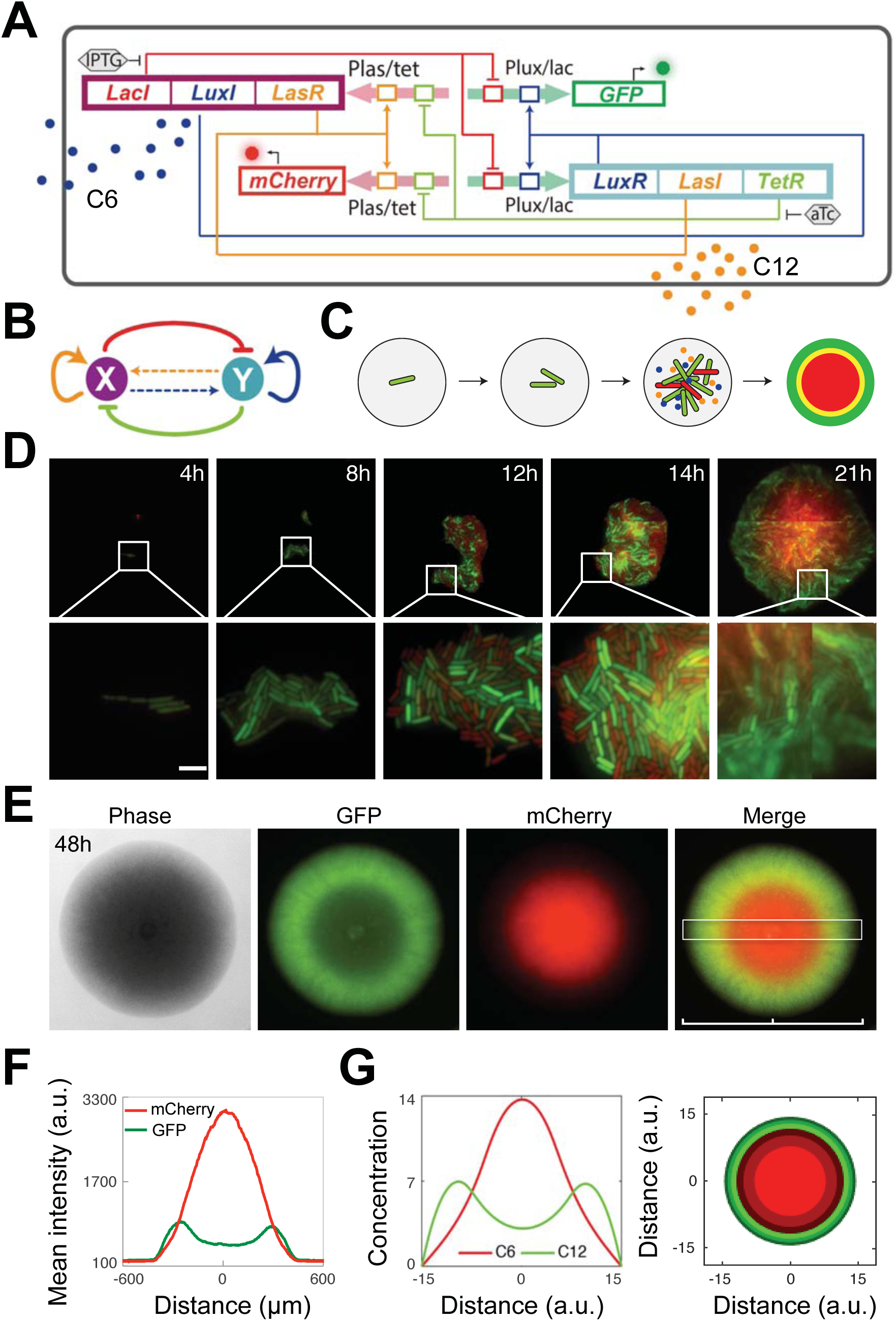
Conceptual and experimental design of MINPAC, and reaction-diffusion based pattern formation. (**A**) Experimental design of the MINPAC network. *Plas/tet* (pink arrow) can be activated by LasR (yellow) and repressed by TetR (light green), while *Plux/lac* (green arrow) can be activated by LuxR (blue) and repressed by LacI (red). LuxI (blue) synthesizes C6 (blue dots) to bind with LuxR to activate *pLux/lac*, while LasI (yellow) synthesizes C12 (yellow dots) to bind with LasR to activate *Plas/tet*. GFP and mCherry serve as reporters for *Plux/lac* and *Plas/tet*. (**B**) Abstract diagram of MINPAC topology, where X and Y mutually inhibit each other (T-bars) and auto-activate (arrowheads) itself, meanwhile X and Y can mutually activate through small autoinducer mediated intercellular communication (dashed arrowheads). Genes and regulations are color-coded corresponding to the circuit in (**A**). (**C**) Illustration of a ring pattern formation from a single *E. coli* cell harboring MINPAC circuit. (**D**) MINPAC directs single cells to self-organize into ring pattern at microscopic scale. Representative experiments of pattern formation from single cell to colony by time-lapse microscopy (Scale bar represents 5 µm). The 21-hr image is captured and combined by four individual images. (**E**) MINPAC cells self-organized double-ring pattern at colony scale. Representative fluorescence images are taken at 48 hr. Magnification: 2x. (**F**) Mean fluorescence intensity across the center of pattern-generating colony (white box in **E**). Distance indicates the size of the colony. (**G**) Left: PDE model simulations of the extracellular C6 and C12 concentrations, which are corresponding to mCherry and GFP intensities, respectively. Right: Two-dimensional ring pattern simulated from the model, with high C6 concentration (red) for cells in the core and high C12 concentration (green) on the edge of the colony, forming a similar double-ring pattern as in (**E**).

Specifically, the MINPAC topology is built upon two hybrid promoters *Plas/tet* and *Plux/lac*, which harbor high nonlinearity and inducibility (Fig. 1A and Fig. S1). *Plas/tet* drives *LasR*, *LuxI* and *LacI* expression, representing the node X in Fig. 1B, whereas *Plux/lac* regulates transcription of *LuxR*, *LasI*, and *TetR*, representing the node Y. LasI and LuxI are synthases that catalyze the synthesis of autoinducer 3-oxo-C12-HSL (C12) and 3-oxo-C6-HSL (C6), respectively. The two small autoinducers can diffuse out of and into cells to mediate cell-cell communication and coordinate population behaviors on a spatial domain. LasR and LuxR activate *Plas/tet* and *Plux/lac* in the presence of C12 and C6, respectively, forming positive autoregulations. IPTG inhibits the repressive effect of LacI on *Plux/lac*, and aTc counteracts TetR inhibition on *Plas/tet*, forming the mutual inhibitions. Green fluorescent protein (GFP) and mCherry protein serve as the corresponding reporters of *Plux/lac* and *Plas/tet* activities in living cells (Fig. 1A).

To investigate whether MINPAC could direct single cells to self-organize into spatial patterns, we transformed the circuit into *E. coli* cells and serially diluted cell cultures into single cells before seeding on a semi-solid minimal M9 medium (Fig. 1C). Using live single-cell time-lapse fluorescence microscopy, we observed the early stage of pattern formation (Fig. 1D). After an initial phase of uniform fluorescence (4 & 8 hours), we observed that cells differentiated into equivalent numbers of green and red fluorescence in a disordered, seemingly-random, spatial distribution (12 & 14 hours). As microcolonies grew to ~100 μm in diameter (between 14 and 21 hours of growth), a red-center green out-circle fluorescence pattern starts to emerge (Fig. 1D). These results illustrate that our engineered pattern formation is scale-dependent at the early stage and the pattern starts to emerge only after cell number reaches a certain threshold. We reason that as the stochastic growth progresses through time, while outcomes of cell-cell variability are hard to predict initially or at microscopic scale, the population starts to synchronize and converge to a collective behavior and become more predictable as time progress or at macroscopic scale.

To further investigate the circuit’s capability in directing pattern formation at macroscopic scale, we carried out long term experiment by culturing single cell initiated colonies on agar plates up to 96 hours. Time-lapse colony imaging results show that the single colony has no obvious pattern at 15 hr and exhibits a weak yellow flat disk, suggesting cells express either GFP or mCherry are distributed without order (Fig. S2). This is consistent with our microscopic observations. After 24 hr, cells in the colony started to differentially and orderly express GFP and mCherry and self-organize into a stable double-ring pattern of an outer GFP-ring and inner mCherry disk at 48 hr (Fig. 1E and S2), with a small temporary yellow ring between these two rings (Fig. S2). The double-ring pattern is stable with time. Fluorescence quantification also confirms higher GFP expression for cells on the edge of the colony and higher mCherry expression for cells in the center (Fig. 1F).

To rule out the possibility that circuit-independent factors such as nutrition or growth are responsible for the pattern, we tested two control circuits: one with GFP and mCherry expressed from constitutive promoters, and the other one with GFP and mCherry expressed from hybrid promoters *Plas/tet* and *Plux/lac*. No obvious ring patterns were observed at 24 or 48 hrs (Fig. S3). Therefore, we conclude that MINPAC circuit is responsible for the self-organized ring pattern in single colonies.

Toward a quantitative and mechanistic understanding of the ring patterning process, we next built a partial differential equation (PDE) model to mathematically describe the production, regulation, transport, and diffusion of autoinducers C6 and C12. LuxI and LasI’s expression in MINPAC governs the synthesis of C6 and C12, which can diffuse out of and back into cells to further regulate the intrinsic transcriptional network MINPAC and determine cells’ fate spatially. Thus, the extracellular C6 and C12 kinetics serve as a predictive snapshot of the spatial pattern and could represent the differential expression of mCherry and GFP, respectively (see Supplemental materials for more details). Fitted with biologically feasible parameters, our model shows the two autoinducers harbor similar dynamics to experimental fluorescence intensities across the colony and can reproduce experimentally observed ring pattern in two-dimensional geometry (Fig. 1G). Such corroboration between the RD-based PDE model and experimental results further verified that observed ring pattern is the result of MINPAC regulated RD process.

To further investigate how MINPAC directs the generation of ring pattern, we carried out deterministic analysis for the reaction term of the RD model (i.e. the ODE part). Time series shows that MINPAC has an oscillating reaction part (Fig. S4A), suggesting the temporal oscillation could drive an organized pattern formation across the expanding colony. From a network topology point of view, MINPAC is composed of two topologically equivalent motifs where a self-activating node activates the other node and it in turn inhibits the self-activating node (Fig. 2A), each forming a robust positive-plus-negative oscillator topology ^27–29^. A fully symmetric MINPAC topology would rapidly go to stable steady states without oscillation, but little asymmetry between the two motifs would lead to a robust oscillation (Fig. S5). Our model-comparison results show that oscillation from one-motif topology is, as previously reported, highly dependent on the strength of its negative feedback (*τ*), which is vital for cyclic gene expression^27,30,31^ (Fig. 2B). However, the two-motif MINPAC harbors a greater robustness and amplitude against parameter perturbations to generate temporal oscillation (Fig. 2C). Such robustness enhances the likelihood of observing our desired phenotypic outputs from the synthetic gene circuit.

**Fig. 2.**
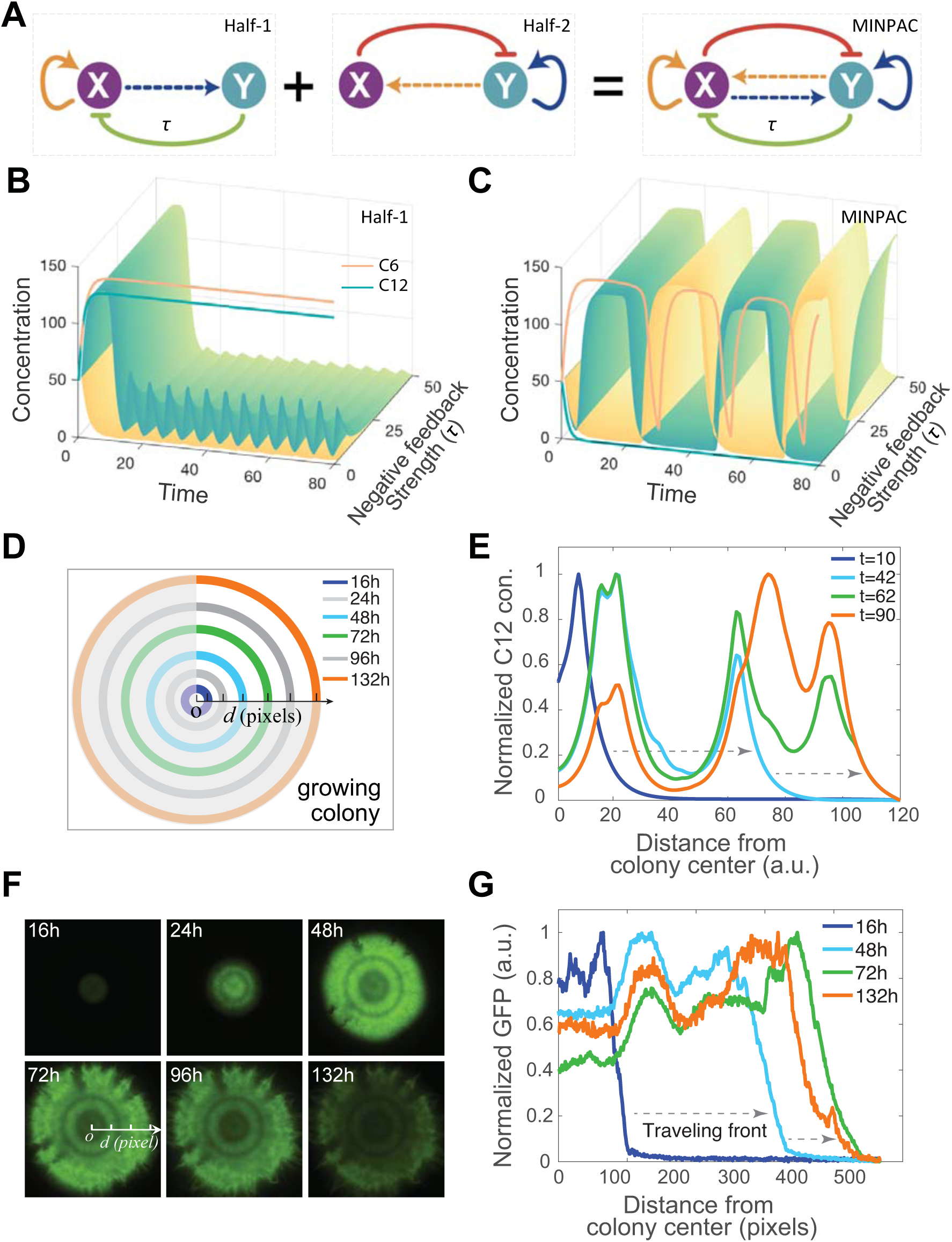
MINPAC directs ring pattern formation through a reaction-diffusion based clock and wavefront mechanism. (**A**) Illustration of the MINPAC composition of two symmetric positive-plus-negative oscillator motifs. Parameter *τ* is used to describe the strength of one negative feedback (node Y inhibits node X). (**B-C**) Model comparison between one-motif topology and two-motif MINPAC. Oscillation from one-motif topology is highly dependent on the parameter *τ* (**B**), whereas MINPAC harbors a greater robustness and amplitude against parameter *τ* changes to generate temporal oscillation (**C**). Cyan and yellow colormaps represent the C6 and C12 concentrations, respectively. The red and blue solid lines are C6 and C12 concentrations when *τ* equals to 0 (i.e. no negative feedback). (**D**) Diagram of a growing colony. Circles with different colors indicate the colony position at different time points. Center is labeled as *o*, and *d* is the distance to the center of the colony. (**E**) Normalized external C12 concentration, directly correlated with experimental GFP intensities, of a pattern-growing colony with time and space from the PDE model simulation. Starting from the center of a colony, colored curves represent C12 concentrations along the colony radius at different time points. Grey arrows indicate the traveling direction of the wave front. (**F**) Time course of a growing colony having multiple GFP rings. (**G**) Quantified temporal and spatial fluorescence intensities of the multiple GFP ring-forming colony in (**F**), showing similar dynamics to model simulation in (**E**). The distance starts from center of the colony from 16 hr to 132 hr. Each pixel is 3.22 μm.

In our MINPAC circuit, promoter functionality tests show LacI is less efficient to inhibit promoter *Plux/lac* (Fig. S1A) compared to tetR to *Plas/tet* (Fig. S1B), supporting that the asymmetric MINPAC could maintain an oscillatory gene expression profile as the molecular clock. Moreover, the autoinducers’ physical diffusion on the agar medium and colony outward expansion (represented as one diffusion term in the PDE model) constitute the propagating wavefront. Finally, the integration of clock and wavefront gates the engineered bacterial cells into subgroups and segment spatially, generating periodic structures. This reaction-diffusion based pattern formation is widely used to explain somitogenesis in development ^2,20,21^.

One interesting phenomenon among vertebrate species is the variations of somite numbers, which is determined by the axis growth and presomitic mesoderm lifetime during embryogenesis ^32,33^. Analogously, we would expect multiple or even indefinite number of stripes for a continuously growing colony (illustrated in Fig. 2D), and colonies with different sizes would have different number of stripes when the oscillation frequency and colony-expanding rates were constant across colonies. With our PDE model, we simulated the temporal dynamics of C12 on the spatial scale and new peaks emerged periodically at the wavefront (Fig. 2E, S4B). Experimentally, ring patterns with multiple stripes were also observed sequentially by time lapse imaging of large colonies (Fig. 2F-G), as model predicted. Collectively, these results suggest that the ring patterns we observed are the outcomes of the spatiotemporal interaction of oscillatory dynamics owing to the network topology and the movement stemming from the diffusion process.

However, even a macroscopic RD system could still be highly sensitive to initial conditions due to the nonlinearity of the network interactions, evidenced by diverse patterns shown in Fig. 3A, some colonies self-organize into a reversed double-ring pattern with GFP accumulating in the inner ring and mCherry on the outer ring (top). A more complicated pattern is also observed, in which two GFP rings alternating with two mCherry rings, forming a multiple GFP-mCherry ring pattern (Fig. 2F and 3A, bottom). Given that these different patterns emerge from the same MINPAC circuit operating in the same cells and under the same conditions, we hypothesize that it is due to random variations of the initial concentrations of intracellular proteins and autoinducers. To computationally test this hypothesis, we tested various initial conditions of the PDE but kept all the parameters the same. The model indeed reproduces the experimental patterns (Fig. 3B). Furthermore, these differences of the patterns suggest the system is not at steady state and, instead, is evolving towards the steady state. The initial condition determines the starting point of the MINPAC system, which will go through a temporal “non-oscillating” spiral (blue line in Fig. 3C) and finally approach oscillation periods (starting from red curve in Fig. 3C). Quantitative simulations show that the oscillatory system, with different initial points, could require significantly different times, so called Poincare return time, to approach the first stable limit cycle (Fig. 3D). Thus, the initial condition and resulting approach-time variances lead to diverse patterns with different stripes (besides colony size). These results illustrate that initial conditions play an important role in shaping the formation of biological patterns, which is consistent with recent theoretical analysis ^16,34^. Furthermore, the experiment-model consistency entices us to use this model to analyze and predict newly emerged patterns under different contexts.

**Fig. 3.**
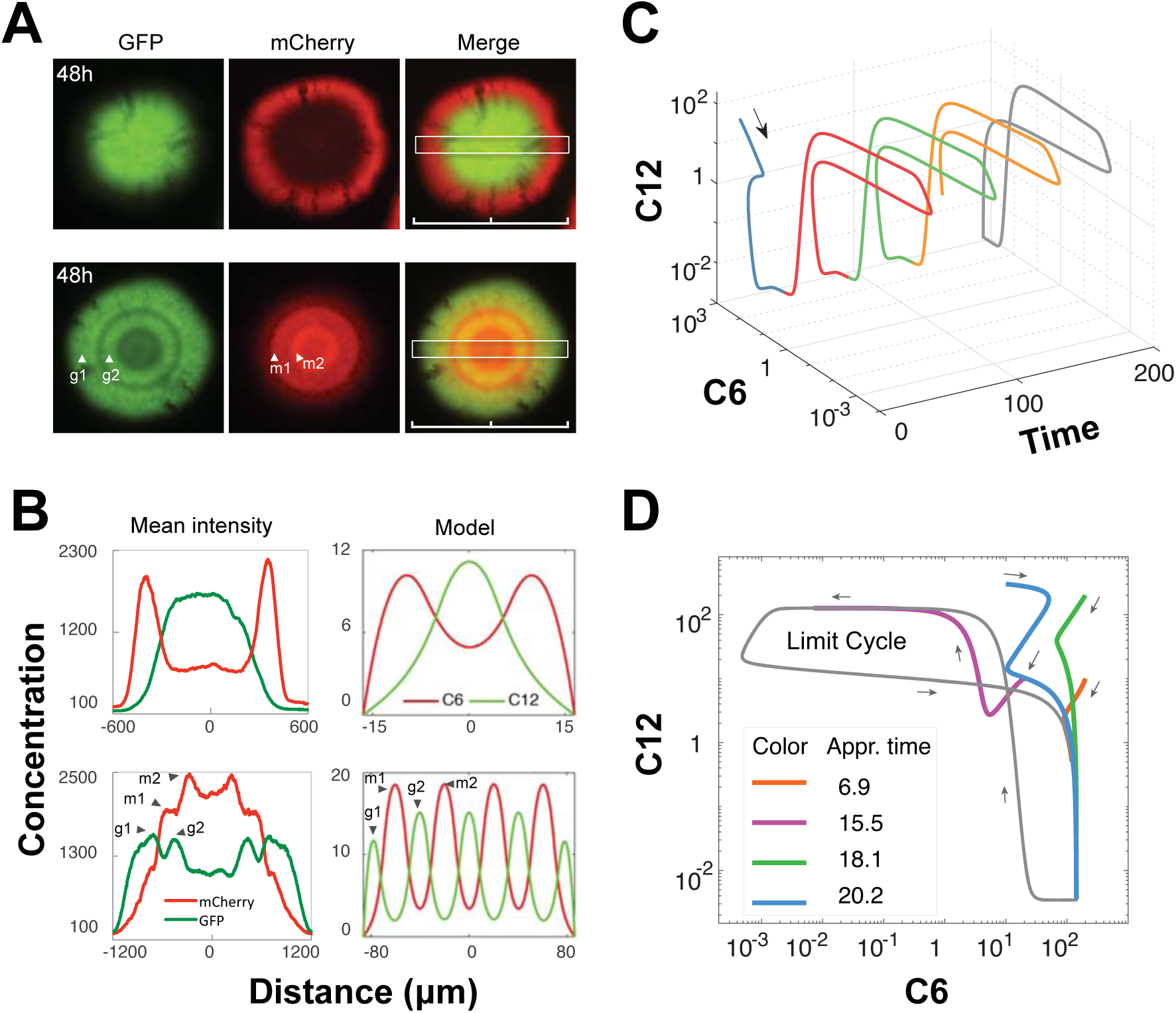
Initial conditions and associated approaching time lead to diverse patterns. (**A**) Two observations distinct to Fig. 1E generated by MINPAC circuit. Top: a ring pattern with a GFP core and a mCherry outer ring; Bottom: a multiple GFP-mCherry ring pattern. (**B**) Left: Mean fluorescence intensities across the center of the ring-forming colonies in (**A**). Rings corresponding to the peaks are labeled. Right: Model simulations recapitulate experimental patterns only through changing the initial conditions of the model. (**C**) A trajectory of a random initial point (black arrow) going to oscillation periods (red, green and yellow curves) simulated from MINPAC reaction term. The grey “butterfly” curve illustrates the limit cycle. (**D**) Approaching time for different initial conditions. Colored curve shows the trajectory before stable oscillations and the approaching time is calculated for the solution going from its starting point to the stable limit cycle (grey curve).

To further examine the pattern’s controllability, we next sought to apply external inducers to perturb the regulations of MINPAC and hence pattern formation. C6, when applied externally, would promote GFP expression and also LasI and TetR production, which could both activate and inhibit mCherry expression. So the net impact of C6 induction is nonlinear and nontrivial. Using the PDE model to simulate C6 application, it is predicted that we can expect a multiple GFP-mCherry ring pattern when MINPAC is induced with external C6 (Fig. 4A, top). Experimentally, we supplemented the medium with 1*10^−8^ M C6 and grow the colony following the same protocol. Results show that the colony first formed an outer GFP ring and a reddish yellow core at 24 hr, which became a red core at 60 hr (Fig. S6A). Strikingly, two GFP rings emerged at 77 hr whereas mCherry mostly accumulated in the center (Fig. 4B top, and Fig. S6A). Quantified fluorescence intensities also illustrate there are four peaks for GFP and one significant peak for mCherry, which is in line with model predictions (Fig. 4B). We noticed the inconsistent dynamics between predicted C6 concentrations and measured mCherry intensities, which is probably because of the slow degradation rate of mCherry protein in living cells. Similarly, external C12 induction results in two GFP rings with unbalanced intensities (Fig. S6B).

**Fig. 4.**
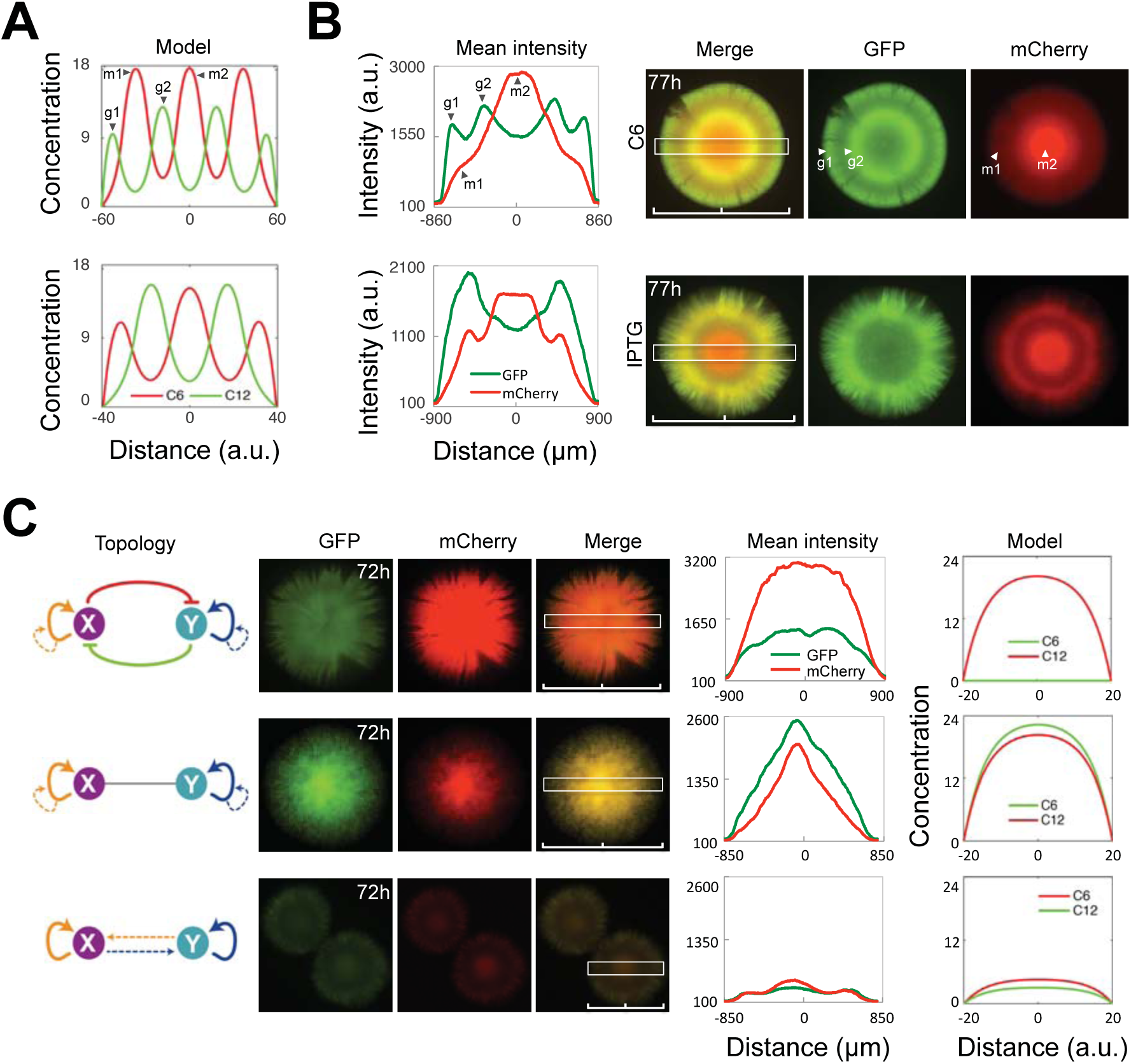
MINPAC directed patterning is tunable and intrinsic to its network topology. (**A**) Model predictions of the pattern under external inducers C6 (top) and IPTG (bottom). (**B**) Experimental validations for model predictions, with C6 and IPTG induction. Top: two GFP rings were observed experimentally under 10^−8^ M C6 induction at 77 hr. Its mean fluorescence intensity across the colony is similar to model prediction (**A**, top). Bottom: a target-like mCherry ring and an outer GFP ring were observed under 10 µM IPTG induction. The mean fluorescence intensity is consistent to model prediction (**A**, bottom). Time course of pattern generation can be found in Fig. S6. (**C**) Three control circuits’ topology and directed patterns. All the circuits are constructed with the same molecular components in MINAPC. Top left: A perturbed MINPAC topology. The intercellular X-Y communications are replaced by intercellular auto-activation of X and Y. No specific pattern is observed experimentally. Middle left: Mutual inhibition is removed and communication is replaced by intercellular auto-activation of X and Y. Strong GFP and mCherry are simultaneously expressed and merged fluorescence is yellow. Bottom left: All regulatory edges are kept but the mutual inhibition module is removed. A weak yellow core and outer ring is observed. Middle: Mean fluorescence intensities across the center of the ring patterns. Right: Model simulations of the three control circuits show consistency to experimental results.

IPTG and aTc induction, on the other hand, can modulate the strength of mutual inhibition in the circuit. IPTG counteracts LacI’s inhibition on *Plux/lac*, leading to more LasI expression and intracellular C12 production. Simulating these changes by perturbing corresponding parameters, the model predicts a target-like mCherry ring with an outer GFP ring pattern (Fig. 4A, bottom), which is further verified by our experimental data (Fig. 4B, bottom). Time course shows that cells in the inner side of the GFP ring started to express mCherry, showing as a yellow ring, at ~60 hr and was stable till 124 hr (Fig. S6A). Inducer aTc’s impacts are similarly predicted and experimentally confirmed (Fig. S6C). Taken together, these results illustrated the controllability of the MINPAC circuit and its directed patterns formation. It is noteworthy that these patterns generated in single colonies autonomously without any predefined spatial cues and the regular structures are robust and stable once formed.

Since the synthetic circuit directed cell-cell communication is established as a viable strategy to generate RD-based and tunable patterns, we employ this method to study the fundamental question of relationship between gene network topology and resulting multicellular pattern. We first designed a perturbed MINPAC topology, where the intercellular X-Y communication modules are replaced by intercellular auto-activations of X and Y (Fig. 4C, top, specific experimental design can be found in Fig. S7). Although there is still autoinducer diffusion, this circuit mitigates the interactions and dependency between X and Y and would remarkably change the intrinsic dynamics. Both experimental observation and model simulation showed no specific pattern but a reddish colony (Fig. 4C, top row). Starting from this topology, we further removed the mutual inhibition module to construct a circuit with two positive feedback motifs (Fig. 4C, middle row), reinforced by intercellular activations. A yellow fluorescent colony with strong GFP and mCherry expression was observed, which is consistent with the model analysis. Lastly, we engineered a sub-network of MINPAC, where the mutual inhibition is removed but keeping the other regulatory edges (Fig. 4C, bottom row). Interestingly, this mutual-activation topology drives a weak yellow target-like ring pattern with low GFP and mCherry expression (Fig. 4C, bottom row). Previous theoretical studies demonstrated that mutual-activation circuit with autoregulations is multistable, and harbors a big parameter space for low-low state ^35,36^. Our model analysis also confirms the low-GFP and low-mCherry expression in this sub-network (Fig. 4C, bottom row). Taken together, each control circuit with different topology has different fluorescence patterns but none of them show the alternating ring patterns, indicating that the multiple-ring pattern is unique to MINPAC circuit.

Biological pattern formation requires complex gene regulation networks and accurate cell-cell coordination. Indeed, coordinated cell population behavior in response to self-regulated morphogen kinetics is a common phenomenon in development ^8,37,38^. Here, we present the design and assembly of a synthetic gene network MINPAC, capable of directing engineered single cells to form self-organized tunable patterns with multiple rings. The PDE model simulations and experimental measurements strongly support that the observed ring patterns are driven by a RD based oscillatory gene network with propagating wavefront, the so-called clock and wavefront mechanism. It is noteworthy to point out that we used one single PDE model to recapitulate and predict all the MINPAC-directed biological patterns. Furthermore, we verified the close connections between gene network topology (circuit architecture) and its induced spatial pattern formation.

MINPAC is a complete motif composed of intracellular transcriptional network and intercellular communication modules, both of which cross-regulate each other to direct spatial pattern formation involving the coordination of molecular gene expression, cellular population response, and positional information interpretation. In this view, the MINPAC represents a paradigm for future design of pattern-forming circuits. Moreover, similar natural counterparts of MINPAC design can be found in the interaction networks of gap genes for the anterior-posterior axis patterning in *Drosophila* ^39–41^. Collectively, this work provides a bottom-up synthetic biology approach to generate complex spatial patterns arising from well-designed reaction-diffusion circuit motif, and integrates experimental data with analytical framework across time and spatial scales to shed lights on the molecular mechanisms of somitogenesis and biological pattern formation, which would contribute to a better understanding of the natural developmental processes, and facilitate the engineering of synthetic tissues in the future.

## Acknowledgments

We thank Dr. James J Collins for the *E. coli* K-12 MG1655 strain with *lac*^−/−^, and Dr. Saeed Tavazoie for lab access for the single-cell microscopy experiments. F.W. was supported by American Heart Association Predoctoral Fellowship 15PRE25710303 (F.W.). This study was financially supported by National Science Foundation Grant DMS-1100309 and National Institutes of Health Grant GM106081 (to X.W.), 5R01GM131405-02 (to Y.K.), and by a NIH Director’s Early Independence Award to K.R.A. (DP5OD019792).

## Author contributions

F.W. and X.W. designed the research. F.W. performed the molecular cloning and patterning experiments. X.F. and K.R.A. designed and performed time-lapse microscopy experiments (agarose pad). T.O., X.C., and Q.Z. participated in the growth condition experiments. F.W., C.H., J.B., Y.K., and X.W. developed the mathematical modeling and computational analysis. F.W., C.H., F.X., Y.K., K.R.A., and X.W. analyzed the data and wrote the manuscript. X.W., K.R.A., and Y.K. supervised the study.

## Data and materials availability

All the experimental materials and procedures and mathematical models are in the supplementary materials. All other data and code are available from the corresponding author upon reasonable request.

## Supplementary information

Materials and Methods

Table S1 – S4

Fig. S1 – S7

References (1 – 15)

