## Supplemental materials for "A Synthetic Biology Approach to Sequential Stripe Patterning and Somitogenesis"

### Supplementary Material

#### 1 Materials and Methods

##### Strains, media and chemicals

All cloning experiments were performed in *Escherichia coli* DH10B (Invitrogen). MINPAC was transformed into *E. coli* K-12 MG1655 strain with *lacI-/-* (*1*) to grow pattern. Cells during cloning were cultured in liquid or solid Luria-Bertani (LB) broth medium with 100 µg/mL ampicillin at 37°C. Pattern grown on minimal salt medium (M9) supplemented with 100 µg/mL ampicillin and 0.45% glucose at 37°C. Chemicals N-(β-Ketocaproyl)-L-homoserine lactone (C6, Sigma-Aldrich), isopropyl β-D-1-thiogalactopyranoside (IPTG, Sigma-Aldrich), and anhydrotetracycline (aTc, Sigma-Aldrich) were dissolved in ddH2O and diluted into indicated working concentrations. N-(3-Oxododecanoyl)-L-homoserine lactone (C12, Sigma-Aldrich) was dissolved in dimethyl sulfoxide (DMSO, Sigma-Aldrich) to avoid precipitation. Liquid cultures were shaken in 15 mL tubes at 220 rotations per minute (rpm).

##### Plasmid construction

Plasmids were constructed using standard molecular biology techniques and all genetic circuits were assembled based on standardized BioBrick methods. Detailed description can be found in reference (*2*). During molecular cloning, both fragment and vector were separated on 1% TAE agarose gel electrophoresis and purified using PureLink gel extraction Kit (Invitrogen). Purified fragment and vector were then ligated by T4 DNA ligase (New England Biolabs, NEB). The ligation products were further transformed into *E. coli* DH10B and plated on LB agar plate with 100 µg/mL ampicillin for screening. Finally, plasmids extracted by GenElute HP MiniPrep Kit (Sigma-Aldrich) were further confirmed through gel electrophoresis (digested with *EcoRI* and *PstI*) and DNA Sequencing (Biodesign sequencing Lab, ASU). All the biological parts used in the paper are obtained from iGEM Registry (http://parts.igem.org/Main_Page) and listed in Table S1. Promoters and each bio-parts were tested before assembly of MINPAC (Fig. S1).

**Experimental set-up for pattern growing and microscopy**

One microliter overnight cultured *E coli* cells harboring MINPAC were serially diluted with M9 medium without antibiotic to $5*10^5$ ~ $5*10^6$ fold and 5 microliter dilution were then evenly plated on the 10 mL semi-solid M9 minimal medium supplemented with 1 mM MgSO4, 100 μM CaCl2, 100 μg/mL ampicillin, 0.01% g/mL amino acid mixtures and 0.45% glucose on a 5-cm petri dish. Fresh M9 solid medium are made for patterning experiment each time. It should be noted that we tried our best to keep the concentration of amino acid mixture around 0.01% g/mL, however, the real concentration is 0.005%~0.02% g/mL because of the limited accuracy of electronic balance and weighting error. For optimal patterning and imaging convenience, we use 0.4% g/mL agarose (colorless), instead of agar (yellow), to solidify the medium and decrease the influence of the background color on microscopic results. After serial dilutions, the cell concentration is ~1 cell (colony) per microliter according to our experience. And the colony size is negatively correlated with the number of the total colonies in the plate, each plate has 0 to 10 separate colonies in our settings. Petri dishes were covered with parafilm (Pentair, US) to decrease the drying process of the medium and grown in 37°C incubator. To make the colonies grow larger, petri dishes were placed in a small cabinet with external water to increase the air humidity. For the induction experiments, we added the inducers (C6, C12, and aTc, and IPTG) to the LB medium (~37°C) before plating into the petri dishes.

Images were taken at indicated times using Nikon Eclipse Ti inverted microscope (Nikon, Japan) at 2x magnification. For continuous observation, the relative position of each colony in the petri dish was labeled and recorded. Exposure time are kept the same during the time course experiments, however, appropriate adjustments are made at some instances especially for the first time point's (12~16 hour) result because of little fluorescent protein expression before 16 hr. We normalized the exposure time from fluorescence intensities to analyze the time course results in Fig. 2F. Brightness and contrast are slightly tuned for presentation of images in the research, however, the mean fluorescence intensities shown in these figures (white boxes on the fluorescence images) are directly acquired from the original images and averaged their intensity values for all the pixels in the white box under the same directions of x-axis (a line goes through the center of the colony). GFP was visualized with an excitation at 472 nm and emission at 520/35 nm and mCherry was visualized with excitation at 562 nm and emission at

641/75 nm using Semrock band-pass filters. For each patterning experiment, we captured ~6 individual colonies and repeated at least two times on different days.

##### Flow cytometry measurements

Flow cytometry measurements were performed using Accuri C6 flow cytometer (Becton Dickinson) and all samples were analyzed at twelve hour and twenty-four hour time points with 488 nm excitation and 530 ± 15 nm emission detection for GFP, and 610 LP for mCherry. For the test of promoter *Plux/lac*, we chose the 12 hrs data to plot, and for *Plas/tet*, we chose the 24 hrs data to plot. 20,000 individual cells were analyzed for each sample at a slow flow rate. Experiments were repeated two times with three replicates. Data files were further analyzed by MATLAB (MathWorks).

##### Time-lapse fluorescence microscopy

*E. coli* K-12 MG1655 strain containing MINPAC was inoculated from frozen stock into growth medium (M9 minimal medium supplemented with 0.1% amino acid, 0.45% glucose and 100 μg/mL Ampicillin). The cells were grown for 12hrs with aeration (300 rpm) at 37°C. To capture colony formation from a single cell, 2% low melting point agarose pad containing 50 μg/mL Ampicillin were prepared according to the previous report (*3*). 2μL diluted overnight culture was put onto agarose pads and slides were incubated at 37°C in dark. Microscopic images were acquired at different time points using Zeiss Axio inverted fluorescence microscope at 63× magnification (Plan-Apochromat 63x/1.4 DIC), with a Zeiss axiocam 503 mono camera. Images covering approximately 1100 μmX800 μm area were collected and processed by Zeiss Zen2 software.

#### 2 Mathematical modeling

##### Model construction

MINPAC circuit is composed of two hybrid promoters (*Plux/lac* and *Plas/tet*), two activators (LuxR and LasR), two repressors (LacI and TetR), two AHL autoinducer synthases (LuxI and LasI), and two promoter reporters (GFP and mCherry). LasI and LuxI are two synthases responsible for the synthesis of autoinducer 3-oxo-C12-HSL

(C12) and 3-oxo-C6-HSL (C6), respectively. The two small autoinducers (morphogens) can diffuse out of and back into cells to mediate cell-cell communication and coordinate population behaviors on a spatial domain. *Plux/lac* activity is determined by the relative concentrations of LacI and LuxR-C6, which is a complex of LuxR protein and intracellular C6 ($C_i$). Similarly, *Plas/tet* dynamics is determined by the relative concentrations of TetR and LasR-C12, which is a complex of LasR protein and intracellular C12 ($H_i$).

To develop a quantitative and mechanistic understanding of the MINPAC-directed ring patterning process, we developed a partial differential equation (PDE) model based on the reaction-diffusion process involving the regulation, production, and diffusion of morphogens C6 and C12. For simplicity, we here used only one equation to overall describe the DNA transcription and protein translation processes, instead of separately describing them. Both the activation and repression are described as hill functions. Based on the biochemical reactions depicted in Fig. 1A, we first derived the ordinary differential equations for LasR ($S$), LacI ($C$), LuxI ($U$), LuxR ($X$), TetR ($E$), and LasI ($A$):

$$\frac{\partial S}{\partial t} = \beta_1 + \frac{k_{11} \cdot (S \cdot H_i)^{n1}}{1 + (S \cdot H_i)^{n1}} \cdot \frac{1}{1 + E^{m1}} - d_{11} \cdot S \quad \text{(Eq1)}$$

$$\frac{\partial C}{\partial t} = \beta_1 + \frac{k_{12} \cdot (S \cdot H_i)^{n1}}{1 + (S \cdot H_i)^{n1}} \cdot \frac{1}{1 + E^{m1}} - d_{12} \cdot C \quad \text{(Eq2)}$$

$$\frac{\partial U}{\partial t} = \beta_1 + \frac{k_{13} \cdot (S \cdot H_i)^{n1}}{1 + (S \cdot H_i)^{n1}} \cdot \frac{1}{1 + E^{m1}} - d_{13} \cdot U \quad \text{(Eq3)}$$

$$\frac{\partial X}{\partial t} = \beta_2 + \frac{k_{21} \cdot (X \cdot C_i)^{n2}}{1 + (X \cdot C_i)^{n2}} \cdot \frac{1}{1 + C^{m2}} - d_{21} \cdot X \quad \text{(Eq4)}$$

$$\frac{\partial E}{\partial t} = \beta_2 + \frac{k_{22} \cdot (X \cdot C_i)^{n2}}{1 + (X \cdot C_i)^{n2}} \cdot \frac{1}{1 + C^{m2}} - d_{22} \cdot E \quad \text{(Eq5)}$$

$$\frac{\partial A}{\partial t} = \beta_2 + \frac{k_{23} \cdot (X \cdot C_i)^{n2}}{1 + (X \cdot C_i)^{n2}} \cdot \frac{1}{1 + C^{m2}} - d_{23} \cdot A \quad \text{(Eq6)}$$

where $\beta_1$ and $\beta_2$ are the basal expressions from *Plas/tet* and *Plux/lac*, respectively. The middle term in the each equation is the positive feedback from LasR-C12 or LuxR-C6 complex and negative feedback from TetR or LacI, respectively. The last term of each equation is the degradation term. $n_1$ and $n_2$, $m_1$ and $m_2$ are the hill coefficients for activation or repression to the promoters from protein complex, like the LuxR-C6 dimer and LacI tetramer. Parameters $k_{ij}$ represent the production rates and $d_{ij}$ represent the degradation rates for proteins LasR, LacI, LuxI, LuxR, TetR, and LasI, respectively.

LasR, LacI, LuxI are produced from the same promoter *Plas/tet* and have similar production terms. We assume the three proteins have similar degradation rates and similar dynamics and use LuxI to represent the LasR and LacI. Similarly, we use LasI to represent LuxR and TetR expression dynamics from *Plux/lac*. Thus the above six equations are simplified by the following two equations:

$$\frac{\partial U}{\partial t} = \beta_1 + \frac{k_1 \cdot (U \cdot H_i)^{n1}}{1 + (U \cdot H_i)^{n1}} \cdot \frac{1}{1 + A^{m1}} - d_1 \cdot U \quad \text{(Eq7)}$$

$$\frac{\partial A}{\partial t} = \beta_2 + \frac{k_2 \cdot (A \cdot C_i)^{n2}}{1 + (A \cdot C_i)^{n2}} \cdot \frac{1}{1 + U^{m2}} - d_2 \cdot A \quad \text{(Eq8)}$$

Next, we consider the dynamics of C6 and C12, whose biosynthesis primarily depends on synthases LuxI and LasI, respectively. Internal C6 and C12 can diffuse out of and into cells, and bacterial cells further respond to the autoinducers when their concentrations exceed a certain threshold. Based on this, we described the internal C6 ($C_i$) and C12 ($H_i$) dynamics with following equations:

$$\frac{\partial C_i}{\partial t} = \frac{k_3 \cdot U^{n3}}{K_c^{n3} + U^{n3}} - d_3 \cdot C_i + D_c \cdot (C_e - C_i) \quad \text{(Eq9)}$$

$$\frac{\partial H_i}{\partial t} = \frac{k_4 \cdot A^{n4}}{K_h^{n4} + A^{n4}} - d_4 \cdot H_i + D_h \cdot (H_e - H_i) \quad \text{(Eq10)}$$

where the first term describes the production of $C_i$ and $H_i$ with the production rate $k_3$ and $k_4$, and $K_c$ and $K_h$ are the half maximal effective concentrations for C6 and C12 productions, respectively. Hill coefficients $n_3$ and $n_4$ indicate the nonlinearity for the intracellular synthesis of C6 and C12 from LuxI and LasI, respectively. The middle term is internal degradation of the two molecules with the rate of $d_7$ and $d_8$, and the last term is the diffusion through the cell membrane (molecule transport), with a diffusion rate $D_c$ and $D_h$. $C_e$ and $H_e$ are the external C6 and C12, which diffuse across the cell colony on the M9 agarose medium. We then used the following two equations to describe $C_e$ and $H_e$ dynamics:

$$\frac{\partial C_e}{\partial t} = -D_c \cdot (C_e - C_i) - d_5 \cdot C_e + D_1 \cdot \frac{\partial^2 C_e}{\partial x^2} \quad \text{(Eq11)}$$

$$\frac{\partial H_e}{\partial t} = -D_h \cdot (H_e - H_i) - d_6 \cdot H_e + D_2 \cdot \frac{\partial^2 H_e}{\partial x^2} \quad \text{(Eq12)}$$

where $d_5$ and $d_6$ are degradation rates of external C6 and C12, and $D_1$ and $D_2$ are the diffusion constants for external C6 and C12 across the colony on the medium, respectively.

It is noteworthy that although *Plux/lac* activity is activated by the complex of LuxR and intracellular C6, the quorum-sensing mechanism is cell population density-dependent. In other words, *Plux/lac* can be activated only when the local environmental C6 reaches to a certain threshold. Thus, it is the external C6 and C12 determine the dynamics of MINPAC as well as the patterning process. Furthermore, since mCherry and GFP are two reporters of the hybrid promoters *Plas/tet* and *Plux/lac*, so we can use external C6 (from *LuxI* gene, $C_e$) and C12 (from *LasI* gene, $H_e$) to equivalently simulate mCherry and GFP dynamics.

Taken together, we derived six equations (Eqs 7–12) to model the MINPAC dynamics, including *LuxI*, *LasI*, $C_i$, $H_i$, $C_e$, and $H_e$. And the extracellular C6 ($C_e$) and C12 ($H_e$) kinetics can be used as a predictive snapshot of the spatial pattern, and represent the differential expression of mCherry and GFP, respectively. This two-component reaction-diffusion equations model was then used to understand MINPAC-directed patterning process and predict its responses to external perturbations.

**Parameter fitting and stochastic simulation**

Based on parameters from previous literatures (*2*, *4–6*) and fitted biologically feasible values, we then numerically solve the PDE model with Pdepe package (Mathworks), which has been used to solve initial-boundary value problems for systems of parabolic and elliptic PDEs in the one spatial variable *x* and time *t*. The model contains two parts, ODE part (Eqs 7-10) and PDE part (Eqs 9-12). We first used ode45 package to solve the

ODE term ($U$, $A$, $C_i$, $H_i$), with defined initial conditions (such as [1, 1, 1, 1]) and given parameter sets. Then we use the solutions of $U$ and $A$ to solve the PDE part ($C_i$, $H_i$, $C_e$, $H_e$) by using the bvp5c package. Here we assume that the range of x-axis is large enough so that all variables are 0 at the boundary, thus the initial condition for the PDE part is [1, 1, 0, 0].

It is well known that the two diffusible signaling molecules with different diffusion constants are one of the fundamental requirements to generate Turing patterns (*7–10*). In our system, we use C6 and C12 to mediate the intercellular communication by diffusing in the colony and semi-solid medium. Previous studies indicate that the diffusion rates for C6 and C12 are very similar (less than 1.5 fold difference) (*11*, *12*) and given that their similar chemical structures (C12 only has six more carbons than C6), we here assume they have the same diffusion coefficient ($D_n$). We also assume that the application of external inducers would influence the original diffusion rates of C6 and C12 going through the cell membrane to some extent (owning to the limitation of cell membrane's molecule transport). For example, C6 addition on the medium leads to a slight decrease of diffusion rate of C12 through cell membrane ($D_h$), and vice versa (Table S3).

To mathematically explain the different ring patterns from the same MINPAC circuit in Fig. 1E and Fig. 3A, we infer that it is likely due to the stochasticity of initial cellular state, which could be attributed to gene expression noise in the cells. The inherently stochastic transcription and translation processes and environmental fluctuations lead to variations of each gene expression, and even distinct phenotypes in single cells (*13*, *14*). In our model simulations, through changing the initial conditions for the four intracellular species *LuxI*, *LasI*, $C_i$ and $H_i$ (Table S2), we can successfully recapitulate experimental results (Fig.1E and Fig. 3A). The domain size ($N$) of each colony is also accordingly set to run the simulation.

For no induction scenarios, we used zero boundary conditions to solve the PDE. To predict the patterning responses under external C6 and C12 inductions, we changed the boundary and initial conditions of the PDE model to mimic such experimental perturbations. However, C6 and C12 application would also change the initial external

C6 and C12 value (i.e. $C_e$ and $H_e$). So we set the boundary condition for $C_e$ and $H_e$ to [1, 0] and [0, 1] under C6 and C12 induction, respectively. In addition, since the external application of C6 and C12 further regulates the intracellular genes expression and MINPAC dynamics, so we also changed the initial values for *LuxI*, *LasI*, $C_i$ and $H_i$ (Table S3) under these two scenarios. Experimental results showed a good match with the model predictions (Fig. 3B and Fig. S6).

The induction of IPTG and aTc, on the other hand, tunes the strength of the mutual inhibition in MINPAC circuit. IPTG application counteracts LacI's inhibition on *Plux/lac*, leading to more LasI expression. By increasing the production rates of LasI ($k_2$) and internal C12 ($k_4$) in the model, we predict a target-like mCherry ring with an outer GFP ring pattern, which is further verified by our experimental data (Fig. 3B). On the contrary, inducer aTc alleviates the repressor TetR's inhibition to *Plas/tet* transcription and promotes mCherry expression. So we decreased the production rates of LuxI ($k_2$) and internal C6 ($k_4$) in the model, and increased the basal expression of *Plas/tet* ($b_1$). The prediction suggests that cells would have a dominant mCherry expression under aTc induction, which is also confirmed by experimental result (Fig. S6C).

Together, our reaction-diffusion model mostly recapitulated experimental results and helps us to understand the MINPAC-directed ring pattern formation process and predict the pattern formation from external perturbations. It is necessary to point out that we here didn't consider the environmental effects including space and nutrition limitations on cell growth and pattern formation, even though it might improve our model prediction efficacy. More detailed mathematical analysis is ongoing to further probe how the environmental factors contribute to pattern formation.

**Model development for the Control circuits**

For the three control circuits of MINPAC in Fig. 4C, we employed a similar strategy to model their molecular interactions since they are engineered with the same biological components. All three control circuits have C6 and C12 intracellular synthesis and extracellular diffusion processes, so the dynamics of $C_i$, $H_i$, $C_e$, and $H_e$ are the same as in the MINPAC model (Eqs 9-12). Thus, we only need to change the reaction equations for LuxI and LasI based on the specific circuit topology.

In the first control, the intercellular X-Y communication modules are replaced by intercellular auto-activations of X and Y (Fig. 4C, top), so we modified the positive feedback terms in the LuxI ($U$) and LasI ($A$) equations:

$$\frac{\partial U}{\partial t} = \beta_1 + \frac{k_1 \cdot (U{\cdot}C_i)^{n1}}{1 + (U{\cdot}C_i)^{n1}} \cdot \frac{1}{1 + A^{m1}} - d_1 \cdot U \qquad \text{(Eq13)}$$

$$\frac{\partial A}{\partial t} = \beta_2 + \frac{k_2 \cdot (A{\cdot}H_i)^{n2}}{1 + (A{\cdot}H_i)^{n2}} \cdot \frac{1}{1 + U^{m2}} - d_2 \cdot A \qquad \text{(Eq14)}$$

In the second control circuit, the mutual inhibition module is removed compared to the first control circuit, namely a circuit with two positive feedback motifs (Fig. 4C, middle). So the model can be written as:

$$\frac{\partial U}{\partial t} = \beta_1 + \frac{k_1 \cdot (U{\cdot}C_i)^{n1}}{1 + (U{\cdot}C_i)^{n1}} - d_1 \cdot U \qquad \text{(Eq15)}$$

$$\frac{\partial A}{\partial t} = \beta_2 + \frac{k_2 \cdot (A{\cdot}H_i)^{n2}}{1 + (A{\cdot}H_i)^{n2}} - d_2 \cdot A \qquad \text{(Eq16)}$$

The third control is a sub-network of MINPAC, where the mutual inhibition is removed but having all the other regulatory edges (Fig. 4C, bottom). So the model can be described as:

$$\frac{\partial U}{\partial t} = \beta_1 + \frac{k_1 \cdot (U{\cdot}H_i)^{n1}}{1 + (U{\cdot}H_i)^{n1}} - d_1 \cdot U \qquad \text{(Eq17)}$$

$$\frac{\partial A}{\partial t} = \beta_2 + \frac{k_2 \cdot (A{\cdot}C_i)^{n2}}{1 + (A{\cdot}C_i)^{n2}} - d_2 \cdot A \qquad \text{(Eq18)}$$

It is worth noting that although the parameter symbols in the three control circuits are the same to MINPAC model, their values may be different, especially for the production rates of LuxI ($k_1$) and LasI ($k_2$) as well as the promoter leakages ($b_1$ and $b_2$) because of the distinct architectures and molecular regulations on the promoters. For example, the basal expression in the second circuit (Control-II) should be larger than in the first circuit (Control-I) and MINPAC circuit owning to a lack of repressors and direct positive autoregulation from LuxR and LuxI. Specific parameters are listed in Table S4.

**Traveling wave solution**

MINPAC is composed of two topologically equivalent motifs where a self-activating node activates the other node and it in turn inhibits the self-activating node (Fig. 2A), each forming a robust positive-plus-negative oscillator topology. However, the two motifs in the MINPAC gene circuit might not be fully balanced. Results in Fig. S1 show that the inhibition efficiency of LacI to promoter *Plux/lac* is lower than TetR to promoter *Plas/tet*. According to our previous gene expression metric in polycistronic circuits (*15*), TetR expression in MINPAC circuit is 32.3% higher than LacI expression when *Plux/lac* and *Plas/tet* have the same production rate. Thus, the two motifs are likely to be unbalanced, indicating MINPAC could maintain a robust positive-plus-negative oscillator topology.

The simulations and experiment results suggest that the ring patterns we observed are the outcomes of the spatiotemporal interaction of oscillatory dynamics owing to the network topology and the movement stemming from the diffusion process. After a short time, the solution of the reaction diffusion system approaches the form of a traveling wave. The traveling wave like solution will move forward at a speed asymptotically constant. If the speed is about one unit of length per unit of time, then the wave front resembles the mirror image of an oscillatory trajectory of the reaction system with a small initial value. A faster wave speed will stretch such oscillatory trajectory while a slower wave speed will compact it. The multiple peaks of such oscillatory trajectories give rise to the observed ring patterns.

First, from the simulation result (the 3-D figure in Fig. S4B) we can observe that the solution approximately takes the form of a traveling wave with a constant wave speed after a period of time (about t=100). Our experiment results provide data for up to 132 hours only (Fig. 2F). As a result, we cannot observe a truly constant speed wave solution. Nevertheless, we can see that a traveling wave like solution emerges as time increases. Second, Fig. 2E and 2G show that the simulation results match experiment observations qualitatively for a set of time points. Thus, the target-like ring patterns result from formation of oscillatory traveling-wave-like solutions in our RD models.

**Legends for supplemental figures**

**Fig. S1. Promoter functionality test.** (**A**) Biological circuit to test hybrid promoter *Plux/lac*. Top: circuit construction. LuxR and LacI are individually expressed from a constitutive promoter (CP) to regulate Plux/lac transcription. GFP is a reporter of *Plux/lac*, and maximum GFP can be achieved with the presence of IPTG and C6. Bottom: experimental data (12 hrs) showing GFP response to doses of C6 and IPTG induction. (**B**) Biological circuit to test promoter *Plas/tet*. Data (24 hrs) shows that *Plas/tet* is only activated in the presence of C12 and aTc. mCherry is a reporter of *Plas/tet* activity. Fluorescence was measured by flow cytometry after adding the inducers. Each data point was averaged from three repeated experiments.

**Fig. S2. Time course for the MINPAC-directed ring pattern formation under no inductions.** Scale bar represents 100 μm. Magnification: 2x.

**Fig. S3. Pattern formation from negative control circuits.** (**A**) Circuit with GFP and mCherry expressed from constitutive promoters. High GFP and mCherry simultaneously expressed and a yellow fluorescence disk is observed at 48 hours. Scar bar: 100 μm. (**B**) Circuit with GFP and mCherry expressed from hybrid promoters *Plux/lac* and *Plas/tet*. No obvious ring patterns were observed at 48 hrs.

**Fig. S4. Dynamic comparison between MINPAC and the three control circuits when diffusion rate is 0.** (**A**) Time series simulation of MINPAC system (Top left) shows a stable periodic oscillation for *LuxI*, *LasI*, $C_i$, and $H_i$. However, the Control circuits go to stable steady states. Multiple parameter values were tested and similar results are produced. (**B**) Model simulation for external C6 and C12 dynamics with time and space from the MINPAC reaction-diffusion model. Simulations start from the center of a colony.

**Fig. S5. MINPAC harbors a great robustness and amplitude against parameter changes to generate temporal oscillation.** (**A**) Abstract diagram of MINPAC topology, which is composed of two symmetric classical positive-plus-negative oscillator topologies. Parameters $k_1$, $k_2$, $k_3$, $k_4$ are production rates for each edge of the MINPAC motif; and $\tau_1$, and $\tau_2$ are the inhibition strength of the two negative feedbacks. (**B**) MINPAC quickly goes to stable steady states when all the corresponding parameters are fully symmetric, i.e. $k_1$ equals to $k_2$; $k_3$ equals to $k_4$, and $\tau_1$ equals to $\tau_2$. Similar applications to all the other parameters. (**C-E**) Oscillation emerged when there is a tiny asymmetry between the two symmetric motifs. Keeping all the other parameters unchanged, MINPAC rapidly goes to robust and stable oscillation periods when $k_1$ increases by 0.8% (i.e. $k_1$ = 1.008 * $k_2$) or $k_3$ increases by 8.75% (i.e. $k_3$ = 1.0875 * $k_4$) or $\tau_1$ increases by 1% (i.e. $\tau_1$ =1.01 * $\tau_2$). This is also applied to the cases when decreasing parameters in a small scale, which is another way to introduce the asymmetry.

**Fig. S6. Tunability of the pattern formation from MINPAC using external inducers.** (**A**) Time course results for the MINPAC directed pattern formation with external C6 (top) and IPTG (bottom) inductions. (**B**) Model prediction (left) of pattern formation under C12 induction and experimental validation (right). Both model and experiments showed that $10^{-9}$ M C12 induction resulted in two GFP rings with unbalanced intensities. (**C**) Model simulation (left) predicts a dominant mCherry expression under aTc induction. Experimental result (right) confirmed model prediction that only mCherry is expressed under 2 ng/ml aTc induction. Mean fluorescence intensities are also showed in the middle panel.

**Fig. S7. Topology and experimental design for the three MINPAC control circuits.** (**A**) A perturbed MINPAC topology. The intercellular X-Y communications are replaced by intercellular auto-activation of X and Y. (**B**) Mutual inhibition is removed, and communication is replaced by intercellular auto-activation of X and Y. (**C**) A MINPAC sub-network, having all regulatory edges of MINPAC, except the mutual inhibition module. The right side are the experimental designs for each control circuit, which is engineered with the same bio-components as in the MINPAC. Genes, promoters, and regulations are color-coded corresponding to the topology on the left side.

**Table S1. Bio-components from Registry of standard biological parts.**

| Biobrick number | Abbreviation in the paper | Description |
|---|---|---|
| BBa_K176009 | CP | Constitutive promoter family member J23107 actual sequence (pCon 0.36) |
| BBa_K182102 | Plux/lac | Hybrid promoter (followed by a RBS) activated by LuxR-C6 complex but inhibited by LacI protein |
| BBa_I14015 | Plas/tet | Hybrid promoter activated by LasR-C12 complex, but inhibited by TetR protein |
| BBa_B0034 | RBS | Ribosome binding site |
| BBa_B0015 | Terminator | Transcriptional terminator (double) |
| BBa_C0062 | LuxR | LuxR repressor/activator |
| BBa_C0179 | LasR | LasR activator |
| BBa_C0161 | LuxI | Autoinducer synthase for 3OC6HSL ( 3-oxohexanoyl-homoserine lactone, C6) from *Vibrio fischeri* |
| BBa_ C0178 | LasI | Autoinducer synthase for 3OC12HSL (N-3-oxododecanoyl homoserine lactone, C12) from *Pseudomonas aeruginosa* |
| BBa_C0012 | LacI | LacI repressor from *E. coli* |
| BBa_C0040 | TetR | Tetracycline repressor from transposon Tn10 |
| BBa_J06702 | mCherry | mCherry (optimized monomeric red fluorescent protein) generator |
| BBa_E0040 | GFP | Green fluorescent protein |

**Table S2. Parameters of the three cases for the MINPAC pattern formation under no inductions.**

| Parameter | Fig.1G | Fig.3B (top) | Fig.3B (bottom) |
|---|---|---|---|
| Initial condition | (1,10,6,10) | (8,4,10,1) | (1,100,60,100) |
| M | 1000 | 1000 | 1000 |
| ex | 31 | 31 | 31 |
| N | 15 | 16.5 | 86.3 |
| d4 | 20 | 20 | 20 |
| d6 | 20 | 20 | 20 |
| Dh | 4 | 4 | 4 |
| Dc | 4 | 4 | 4 |
| Dn | 800 | 800 | 800 |
| k1 | 640 | 640 | 640 |
| k2 | 700 | 700 | 700 |
| k3 | 80 | 80 | 80 |
| k4 | 105 | 105 | 105 |
| k5 | 1 | 1 | 1 |
| k6 | 1 | 1 | 1 |
| Kc | 70 | 70 | 70 |
| Kh | 82 | 82 | 82 |
| m1 | 4 | 4 | 4 |
| m2 | 4 | 4 | 4 |
| b1 | 0.8 | 0.8 | 0.8 |
| b2 | 0.5 | 0.5 | 0.5 |
| d1 | 1.19 | 1.19 | 1.19 |
| d2 | 1.19 | 1.19 | 1.19 |
| d3 | 0.56 | 0.56 | 0.56 |
| d5 | 0.8 | 0.8 | 0.8 |
| n1 | 2 | 2 | 2 |
| n2 | 4 | 4 | 4 |
| n3 | 3 | 3 | 3 |
| n4 | 2 | 2 | 2 |

**Table S3. Parameters of the MINPAC prediction under four different inducers.**

| **Parameter** | **C6 induction** (Fig.4A-Top) | **IPTG induction** (Fig.4A-Bottom) | **C12 induction** (Fig. S6B) | **aTc induction** (Fig. S6C) |
|---|---|---|---|---|
| Initial condition | (1,1,20,1) | (2,1,1,1) | (1,1,2,18) | (1,1,1,1) |
| M | 1000 | 1000 | 1000 | 1000 |
| ex | 31 | 31 | 31 | 31 |
| N | 62 | 39.5 | 58 | 19 |
| d4 | 20 | 20 | 20 | 20 |
| d6 | 20 | 20 | 20 | 20 |
| Dh | 3.5 | 3.5 | 4 | 3.5 |
| Dc | 4 | 3.5 | 3.5 | 3.5 |
| Dn | 800 | 800 | 800 | 800 |
| k1 | 640 | 640 | 640 | 640 |
| k2 | 700 | 850 | 700 | 300 |
| k3 | 80 | 80 | 80 | 80 |
| k4 | 105 | 128 | 105 | 80 |
| k5 | 1 | 1 | 1 | 1 |
| k6 | 1 | 1 | 1 | 1 |
| Kc | 70 | 70 | 70 | 70 |
| Kh | 82 | 82 | 82 | 82 |
| m1 | 4 | 4 | 4 | 4 |
| m2 | 4 | 4 | 4 | 4 |
| b1 | 0.8 | 0.4 | 0.8 | 2.6 |
| b2 | 0.5 | 0.45 | 0.5 | 0.5 |
| d1 | 1.19 | 1.19 | 1.19 | 1.19 |
| d2 | 1.19 | 1.19 | 1.19 | 1.19 |
| d3 | 0.56 | 0.56 | 0.56 | 0.56 |
| d5 | 0.8 | 0.8 | 0.8 | 0.8 |
| n1 | 2 | 2 | 2 | 2 |
| n2 | 4 | 4 | 4 | 4 |
| n3 | 3 | 3 | 3 | 3 |
| n4 | 2 | 2 | 2 | 2 |

**Table S4. Parameters of the three control circuits.**

| Parameter | Control I (Fig. 4C, Top) | Control II (Fig. 4C, Middle) | Control III (Fig. 4C, Bottom) |
|---|---|---|---|
| Initial condition | (1,1,1,1) | (1,1,1,1) | (1,1,1,1) |
| M | 1000 | 1000 | 1000 |
| ex | 31 | 31 | 31 |
| N | 20 | 20 | 20 |
| d4 | 20 | 20 | 20 |
| d6 | 20 | 20 | 20 |
| Dh | 4 | 4 | 4 |
| Dc | 4 | 4 | 4 |
| Dn | 800 | 800 | 800 |
| k1 | 740 | 800 | 52 |
| k2 | 800 | 980 | 40 |
| k3 | 80 | 80 | 80 |
| k4 | 105 | 105 | 105 |
| k5 | 1 | 1 | 1 |
| k6 | 1 | 1 | 1 |
| Kc | 70 | 70 | 70 |
| Kh | 82 | 82 | 82 |
| m1 | 4 | - | - |
| m2 | 4 | - | - |
| b1 | 0.5 | 2.5 | 1 |
| b2 | 0.8 | 2.5 | 0.7 |
| d1 | 1.19 | 1.19 | 1.19 |
| d2 | 1.19 | 1.19 | 1.19 |
| d3 | 0.56 | 0.56 | 0.56 |
| d5 | 0.8 | 0.8 | 0.8 |
| n1 | 2 | 2 | 2 |
| n2 | 4 | 4 | 4 |
| n3 | 3 | 3 | 3 |
| n4 | 2 | 2 | 2 |

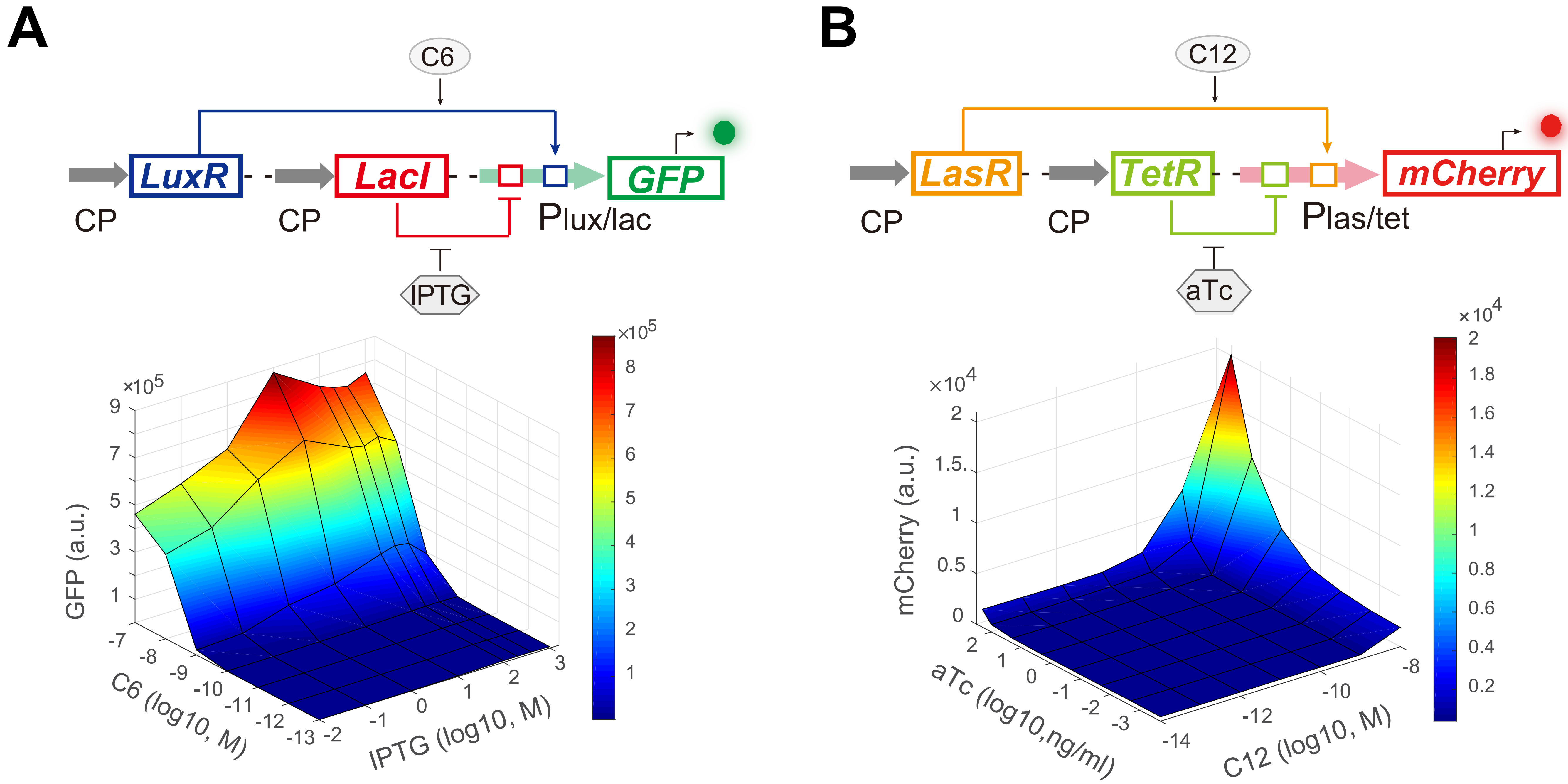

A
C6
LuxR
LacI
GFP
CP
CP
Plux/lac
IPTG
×10^5
GFP (a.u.)
C6 (log10, M)
IPTG (log10, M)
B
C12
LasR
TetR
mCherry
CP
CP
Plas/tet
aTc
×10^4
mCherry (a.u.)
aTc (log10,ng/ml)
C12 (log10, M)


**Fig. S1**

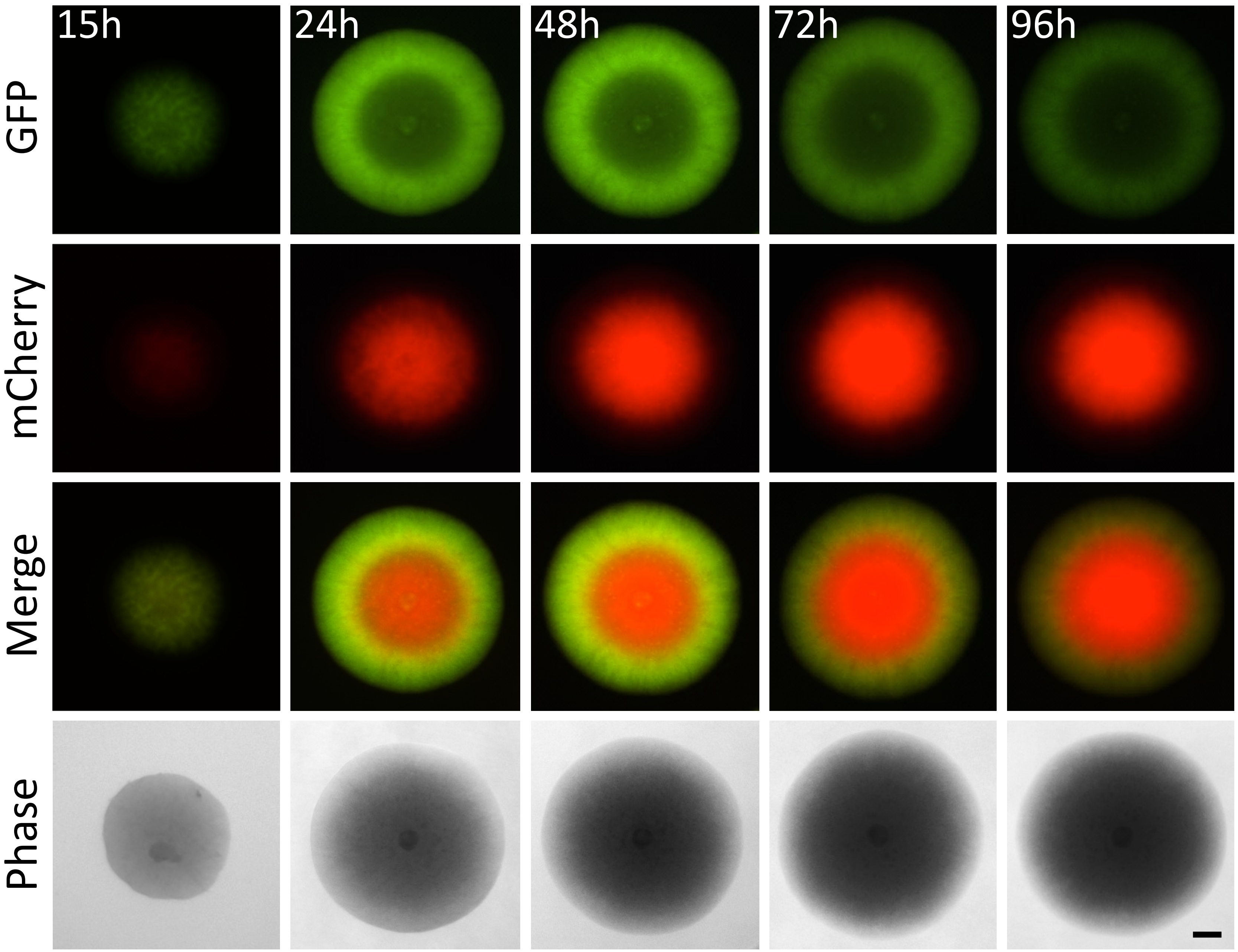


**Fig. S2**

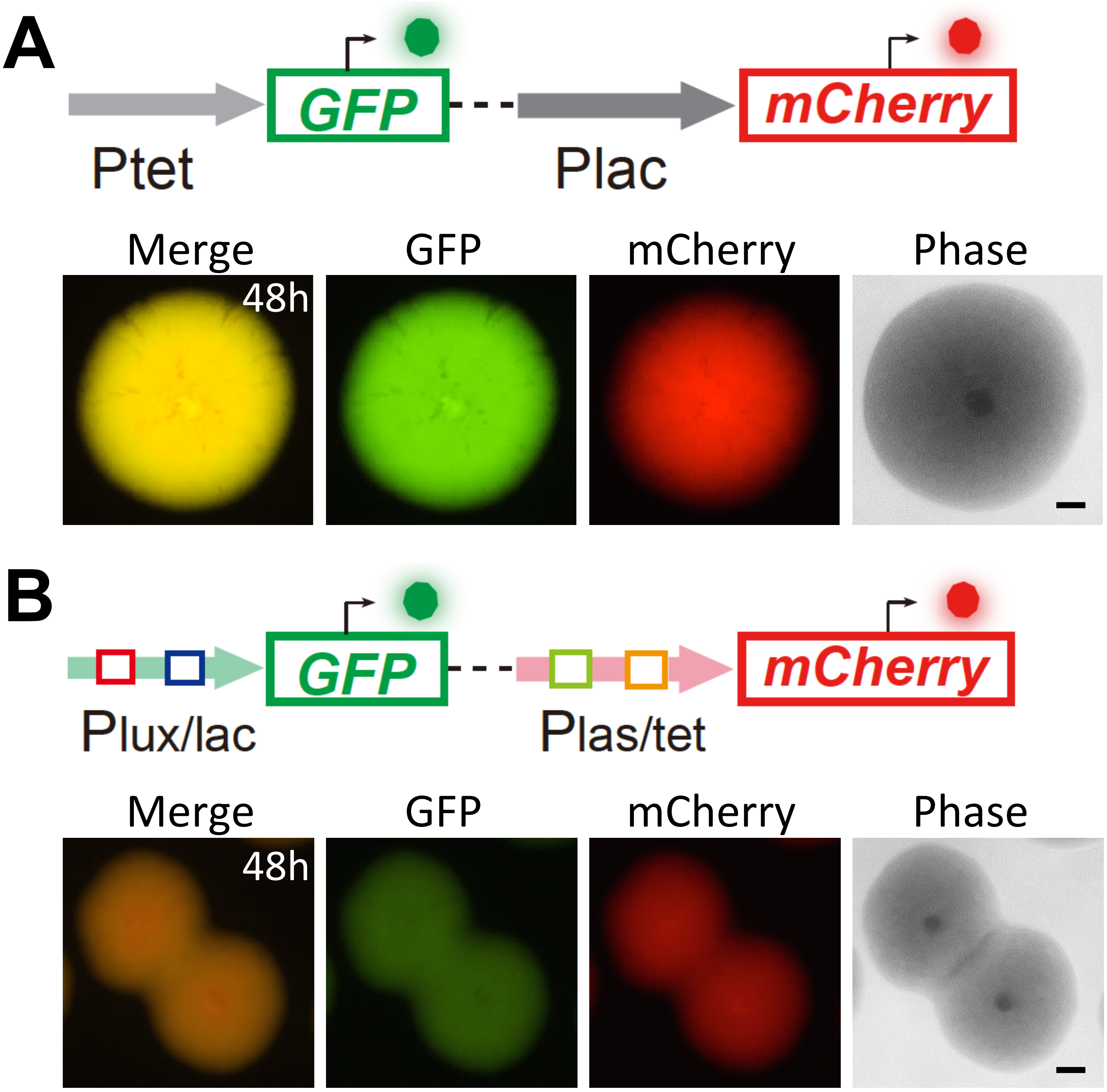
A
GFP
mCherry
Ptet
Plac
Merge
GFP
mCherry
Phase
48h
B
GFP
mCherry
Plux/lac
Plas/tet
Merge
GFP
mCherry
Phase
48h

**Fig. S3**

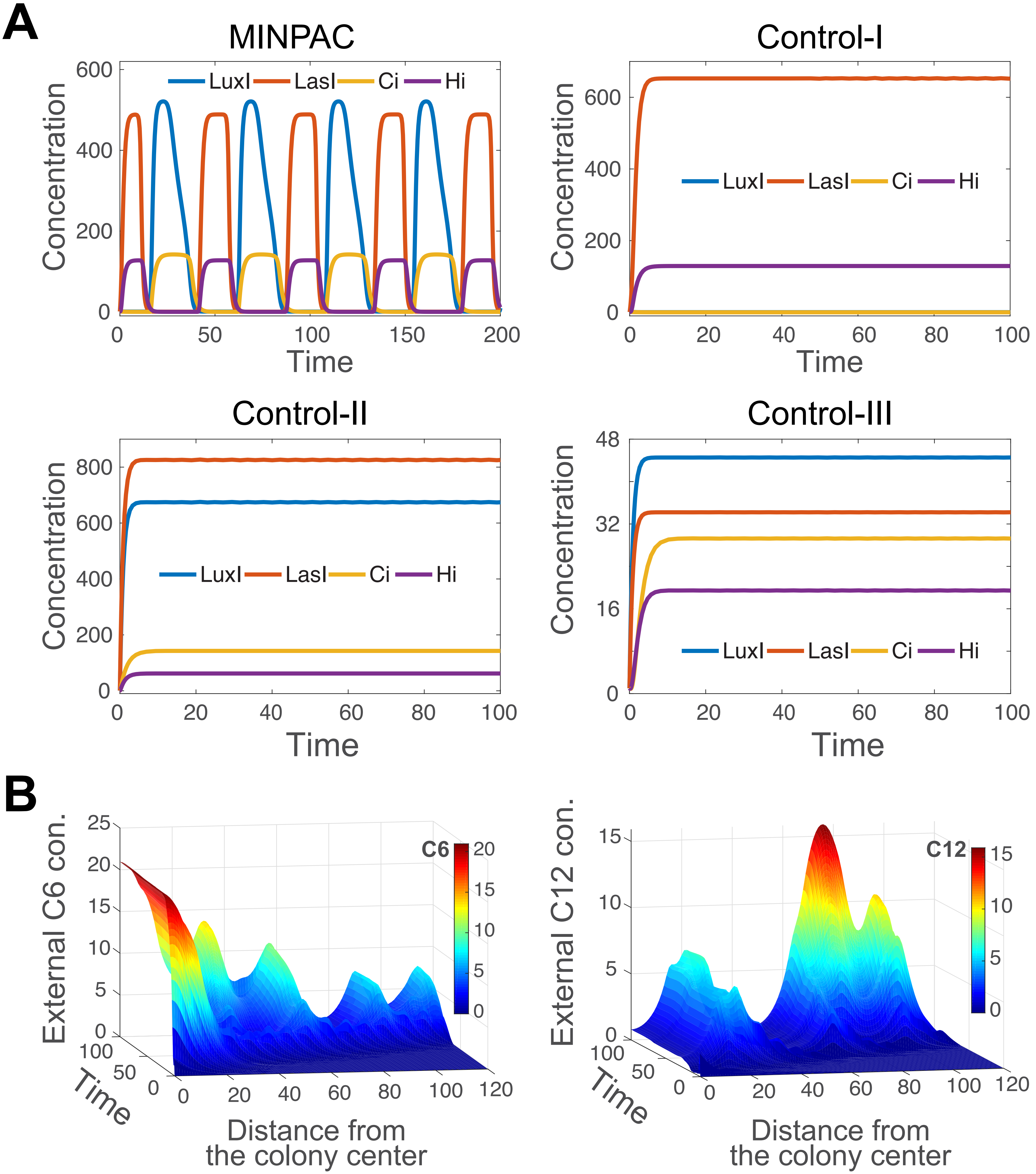
A
MINPAC
Control-I
Control-II
Control-III
LuxI
LasI
Ci
Hi
Concentration
Time
B
External C6 con.
External C12 con.
C6
C12
Time
Distance from the colony center

**Fig. S4**

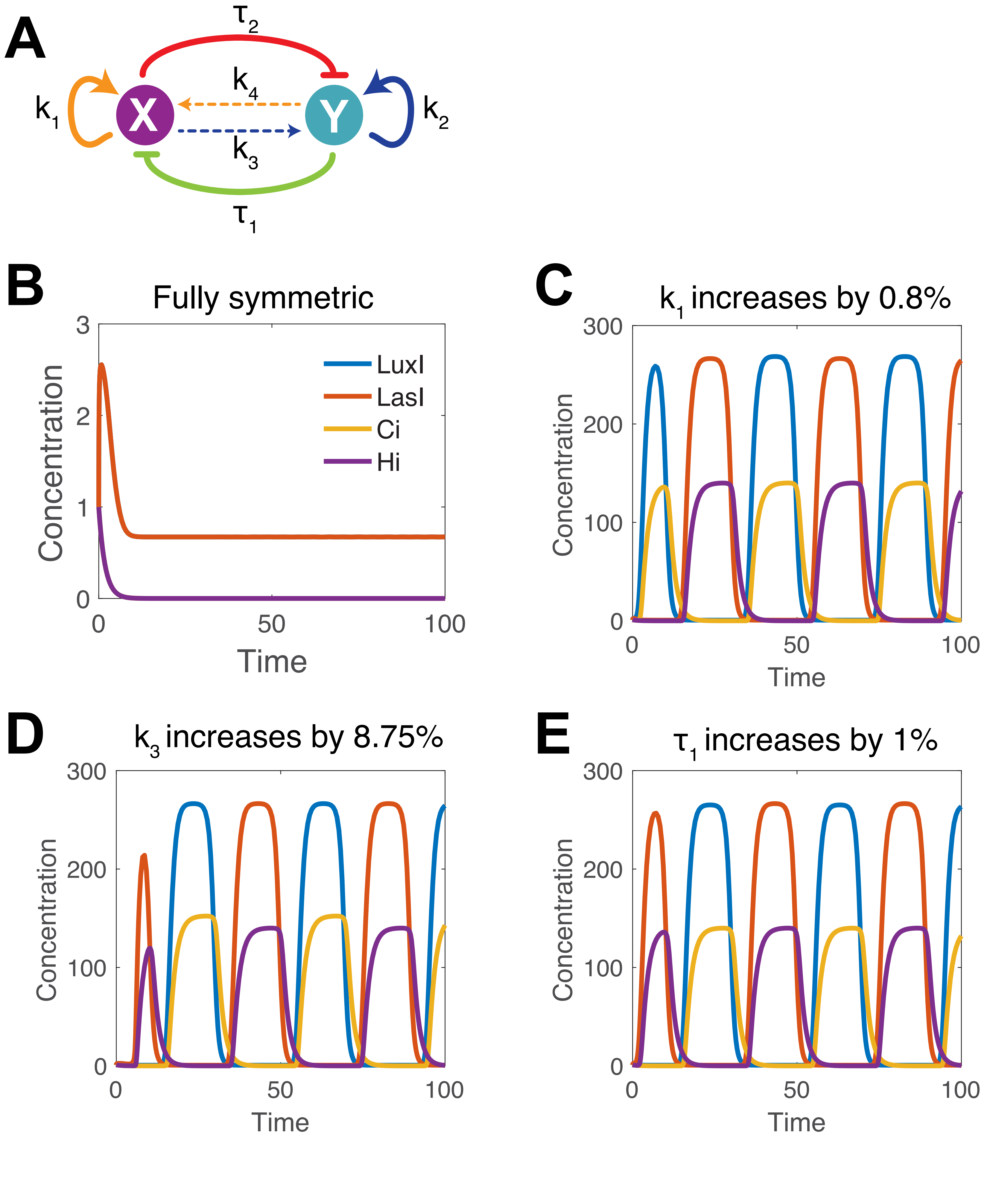
A
B
C
D
E
X
Y
k1
k2
k3
k4
τ1
τ2
Fully symmetric
k1 increases by 0.8%
k3 increases by 8.75%
τ1 increases by 1%
LuxI
LasI
Ci
Hi
Concentration
Time
0
1
2
3
50
100
200
300

**Fig. S5**

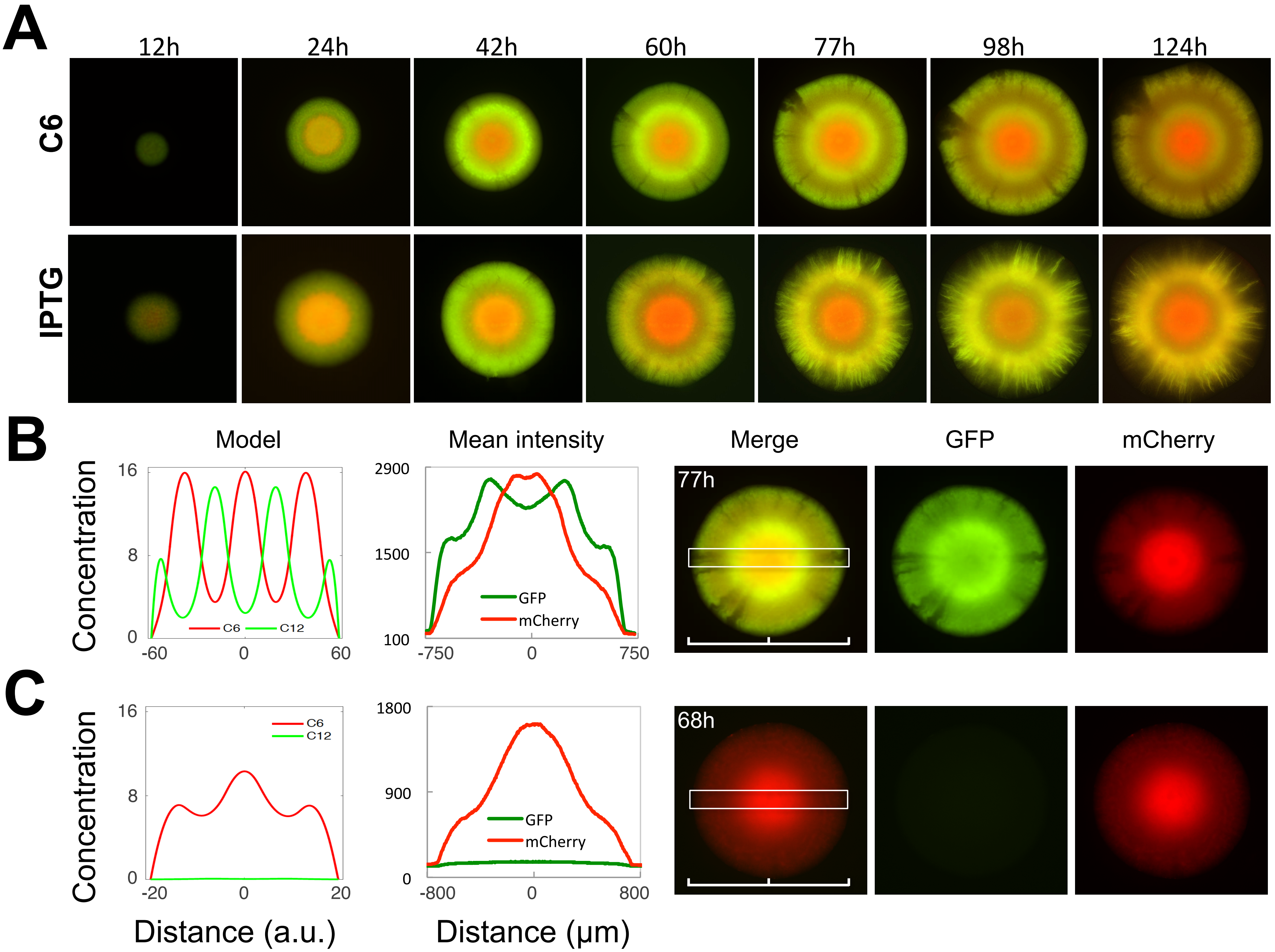
A
12h
24h
42h
60h
77h
98h
124h
C6
IPTG
B
Model
Mean intensity
Merge
GFP
mCherry
Concentration
16
8
0
-60
0
60
C6
C12
2900
1500
100
-750
0
750
GFP
mCherry
77h
C
16
8
0
-20
0
20
1800
900
0
-800
0
800
68h
Distance (a.u.)
Distance (μm)

**Fig. S6**

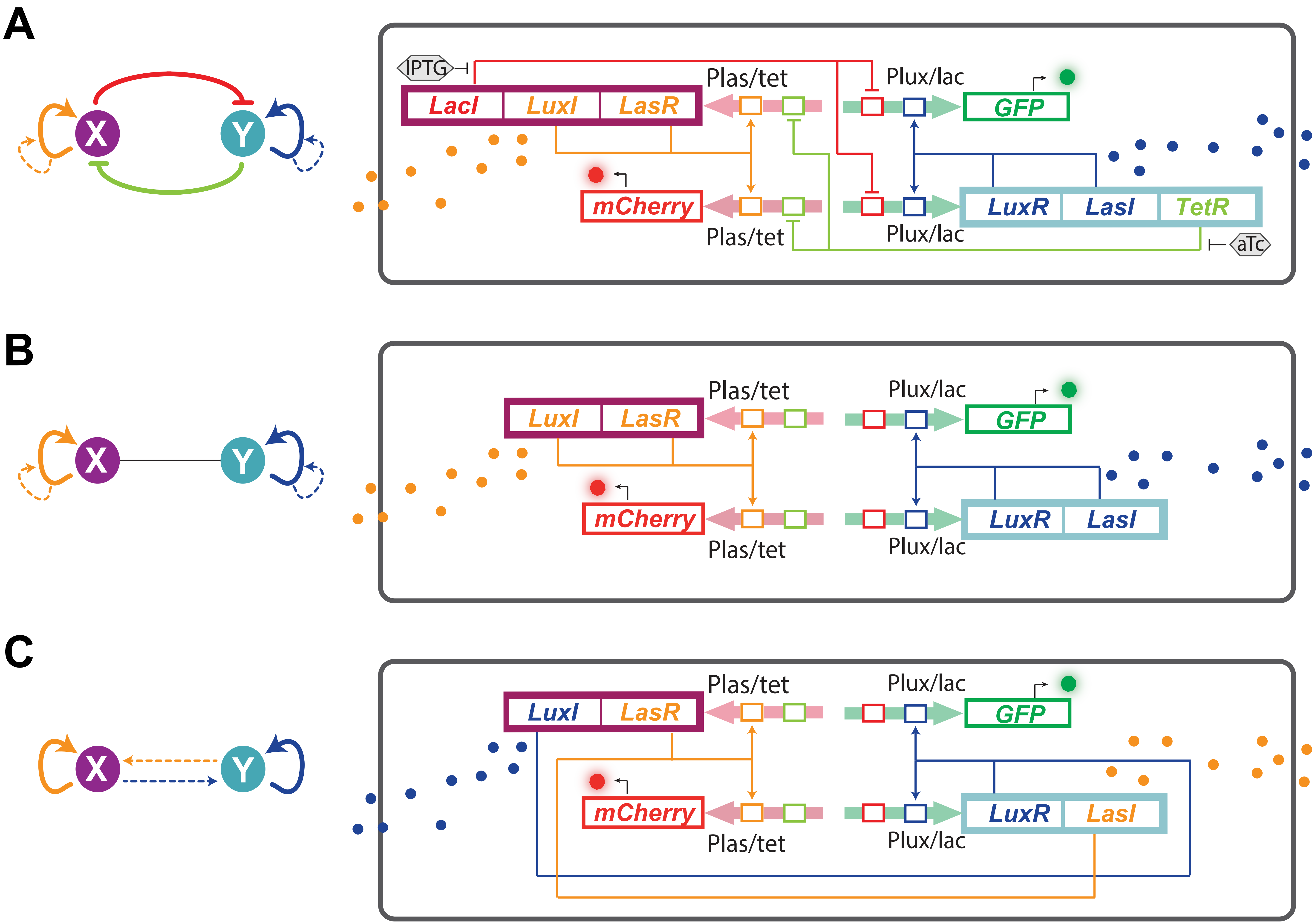
A
X
Y
IPTG
LacI
LuxI
LasR
Plas/tet
Plux/lac
GFP
mCherry
LuxR
LasI
TetR
aTc
B
X
Y
LuxI
LasR
Plas/tet
Plux/lac
GFP
mCherry
LuxR
LasI
C
X
Y
LuxI
LasR
Plas/tet
Plux/lac
GFP
mCherry
LuxR
LasI

**Fig. S7**
